# Damage-induced IL-18 stimulates thymic NK Cells limiting endogenous tissue regeneration

**DOI:** 10.1101/2024.09.27.615528

**Authors:** David Granadier, Kirsten Cooper, Anastasia Kousa, Dante Acenas, Andri Lemarquis, Vanessa Hernandez, Makya Warren, Lorenzo Iovino, Paul deRoos, Emma E. Lederer, Steve Shannon-Sevillano, Sinéad Kinsella, Cindy Evandy, Marcel R.M. van den Brink, Jarrod A. Dudakov

**Author notes:** Corresponding authors: David Granadier, Jarrod A. Dudakov.

## Abstract

Interleukin-18 is an acute phase pro-inflammatory molecule crucial for mediating viral clearance by activating Th1 CD4^+^, cytotoxic CD8^+^ T, and NK cells. Here, we show that mature IL-18 is generated in the thymus following numerous distinct forms of tissue damage, all of which cause caspase-1-mediated immunogenic cell death. We report that IL-18 stimulated cytotoxic NK cells limit endogenous thymic regeneration, a critical process that ensures restoration of immune competence after acute insults like stress, infection, chemotherapy, and radiation. NK cells suppressed thymus recovery by aberrantly targeting thymic epithelial cells (TECs), which act as the master regulators of organ function and regeneration. Together these studies reveal a novel pathway regulating tissue regeneration in the thymus and offer IL-18 as a potential therapeutic target to boost thymic function. Moreover, given the enthusiasm for IL-18 as a cancer immunotherapy for its capacity to elicit a type-1 immune response, these findings also offer insight into potential off-target effects.

## INTRODUCTION

Despite its importance for the production of a diverse and tolerant T cell repertoire, the thymus is exquisitely sensitive to acute insult such as stress-induced rises in corticosteroids and infection, as well as more profound injuries such as chemotherapy and myeloablative conditioning pre-hematopoietic cell transplantation (HCT)^1, 2, 3^. The thymus also harbors an endogenous capacity for regeneration following such acute injuries, however, tissue recovery is a prolonged process such that patients receiving intense thymus-damaging treatments may experience long periods of lymphopenia^4^. This is especially pertinent in HCT-recipients who are particularly vulnerable to opportunistic infection and malignant relapse – two leading causes of post-transplant mortality that are directly related to immune reconstitution – during a months to years-long duration of T cell immunodeficiency^5, 6, 7^. Therefore, understanding the mechanisms underlying thymus recovery could inform therapeutic targets to improve T cell reconstitution in such settings of organ damage ^1^. We and others have identified several cells and molecules that mediate endogenous thymic regeneration including innate lymphoid cells, endothelial cells, eosinophils, and thymic epithelial cells (TECs) and the production of regeneration factors like IL- 22, BMP4, and KGF^8, 9, 10, 11^. However, despite these findings, there remains no clinically approved strategy for treating T cell lymphopenia.

We have previously reported that HCT-conditioning leads to an acute rise in not only apoptosis, but also pyroptosis – a form of immunogenic cell death^12, 13, 14^. During pyroptosis, intracellular contents are released causing local inflammation. We have recently reported that this immunogenic cell death leads to release of damage-associated molecular patterns (DAMPs) such as zinc and ATP that trigger regenerative responses via production of BMP4 by endothelial cells and induction of *FoxN1* expression within thymic epithelial cells (TECs), respectively^15, 16^. However, pyroptosis also leads to the release of inflammatory cytokine IL-18 into the extracellular milieu^12^. IL-18 is a potent stimulator of type II interferon and cytotoxicity profiles in NK cells^17, 18^. NK cells can regulate tissue regeneration and wound healing and recent works implicate NK cells in delaying epithelial recovery^19, 20, 21^. Here we interrogate the impact of this pro-inflammatory cascade and identified that acute thymus damage induced release of IL-18 suppressed endogenous mechanisms of organ recovery by stimulating resident cytotoxic NK cells that aberrantly target TECs.

Overall, this study demonstrates a novel pathway implicating injury induced local inflammation in the suppression of thymic function and isolates the cytokine IL-18 as a critical regulator of endogenous immune reconstitution.

## RESULTS

### Acute thymus injury leads to Caspase-1 cleavage and release of active IL-1β and IL-18

As part of normal T cell development, CD4^+^ CD8^+^ Double Positive (DP) Thymocytes and CD4^+^/CD8^+^ Single Positive Thymocytes undergo apoptosis as newly formed T cell receptors (TCRs) are tested against self-peptide:MHC complexes in positive and negative selection processes ^22^. In fact, approximately 98% of all developing thymocytes fail to pass and survive thymic selection ^23^. Importantly, apoptosis is an immunologically silent process and there is little inflammation within the homeostatic thymus^13^. Following acute damage such as that caused by the cytoreductive conditioning received prior to HCT, modeled with sublethal total body irradiation (SL-TBI, 550 cGy), thymus cellularity precipitously declines^8, 24^ **(Fig 1a)**. However, we have recently reported that ionizing radiation damage leads to cell death by both apoptosis and pyroptosis within the thymus ^16^. In contrast to the immunologically silent apoptosis, pyroptosis is an immunogenic from of cell death mediated by cleaved Caspase 1 (cl-Cas-1) resulting in the release of damage-associated molecular patterns (DAMPs)^12^. Activation of Caspase-1 occurred not only following ionizing radiation, but also following all other forms of acute thymus injuries tested: corticosteroid-induced stress, cytoreductive chemotherapy, and LPS **(Fig. 1a, S1a)**, all of which have been shown to induce acute thymic involution^3, 4, 25, 26, 27^. Although our studies have shown that release of DAMPs such as ATP and zinc can actually be pro-regenerative^15, 16^, cl-Cas-1 also mediates the proteolytic cleavage of the immature, inactive forms of IL-1β and IL- 18 into their mature, active, inflammatory states. Accordingly, cleavage of caspase Cas-1 led to the activation of IL-1β and IL-18 into their mature states within the thymus following each of these acute damage models **(Fig 1b),** and we could detect no increase in activated IL-18 after TBI in mice lacking the catalytic domain of Cas-1 (**Fig. 1c**). IL-18 Binding Protein (IL-18BP), an endogenous antagonist of IL-18, was also upregulated following HCT-conditioning (**Fig. 1d)**; possibly in response to upregulation of activated IL-18^28^. However, while early upregulation of IL-18BP reduced the ratio of IL-18 to IL-18 BP, this was rapidly reversed soon after allowing increased free IL-18 to mediate its pro-inflammatory effects (**Fig. 1e)**.

**Figure 1:**
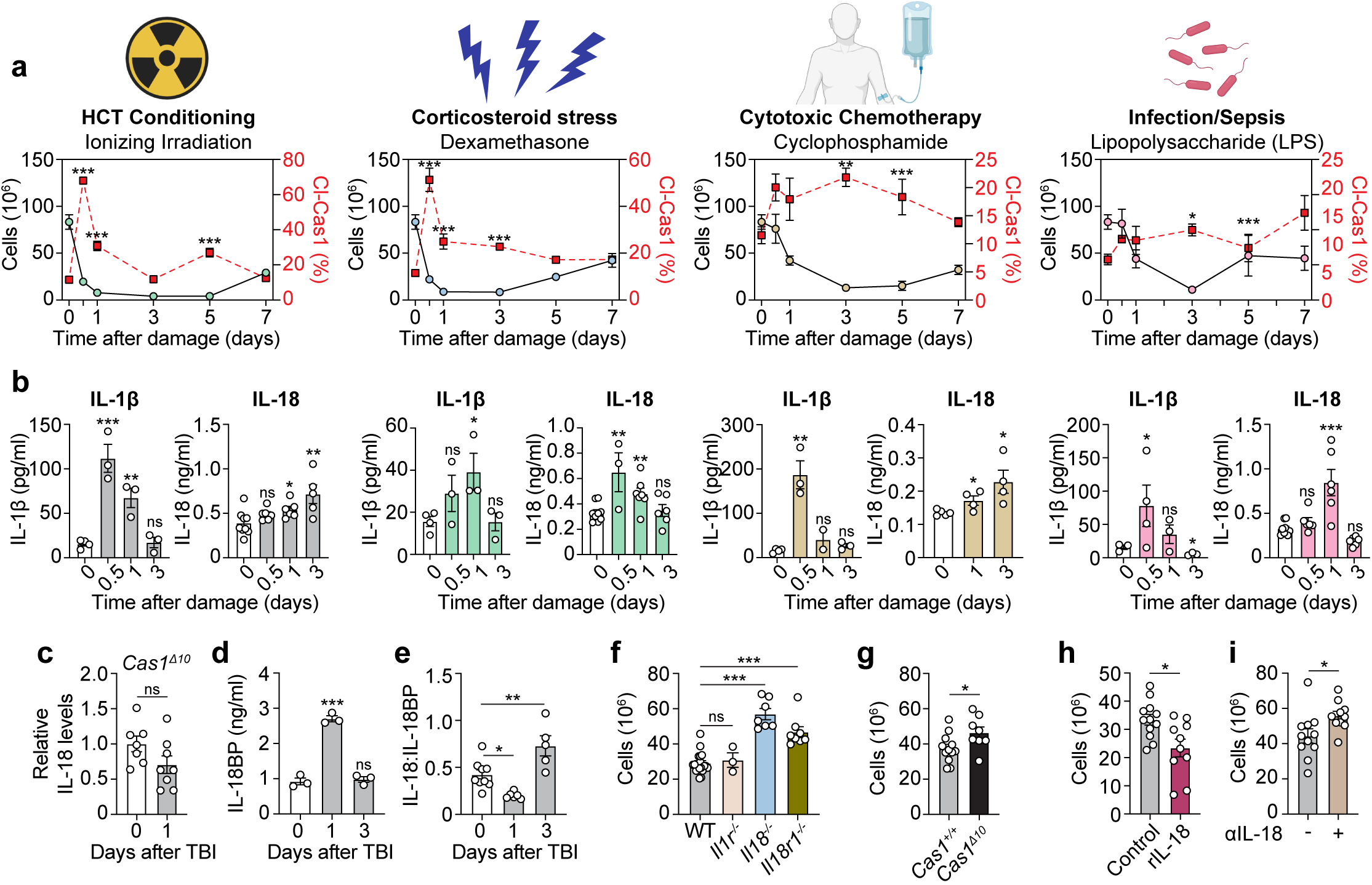
Acute thymic damage triggers cleavage of caspase 1 and activation of IL-18 which suppresses thymus regeneration. **a-c,** Female 1-2 mo C57/BL6 mice were given sublethal total body irradiation (SL-TBI, 550 cGy), Dexamethasone (i.p., 20 mg/kg), Cyclophosphamide (i.p., 200 mg/kg), or LPS (i.p., 1.5mg/kg). **a,** Thymus cellularity (Black) and Cleaved Cas-1 (Red) was measured by fluorescently conjugated FAM-YVAD-FMK at the given timepoints post-treatment (n=4- 15/group/timepoint); all statistics compared to Day 0. **b,** Thymic amount of active IL-1β and active IL-18 (n=3-8/group) measured by ELISA at the indicated timepoints; all statistics compared to Day 0. **c,** Amount of active IL-18 measured by ELISA in female 1-2 mo C57/BL6 WT or *Cas1*^Δ10^ mice at day 1 after SL-TBI (n=6-7/group). **d,** Amount of IL-18 binding protein (IL- 18BP) at days 0, 1 and 3 after SL-TBI (n=3/group). **e,** Ratio of active IL-18 to IL-18BP average at days 0, 1 and 3 after SL-TBI representing free active IL-18 (n=5-8/group). **f,** Female 1-2 mo C57/BL6 WT, *Il1r1^-/-^*, *Il18^-/-^* and *Il18r1^-/-^* mice were given SL-TBI and thymus cellularity was measured 7 days later (n=3-23/group/genotype). **g,** Female 1-2 mo C57/BL6 WT or *Cas1*^Δ10^ mice were given SL-TBI and thymus cellularity measured at day 7 (n=5-10/group). **h,** Female 1- 2 mo C57/BL6 WT mice were given SL-TBI and then administered recombinant IL-18 (rIL-18) on day 3 (s.c., 2.5 mg/kg). On day 7 thymus cellularity was measured (n=10-12/group). **i,** Female 1-2 mo C57BL/6 mice were lethally irradiated and transplanted (i.v.) with 5×10^6^ CD45.1^+^ WT bone marrow hematopoietic cells. Recipient mice were treated with 200μg αIL-18 monoclonal antibody or equal volume control PBS −1, 1, 3, 6, 9, 12, 15 and 18 days and thymus cellularity measured at day 50 post-transplant (n=10-11/group). Graphs represent mean ± SEM. *p<0.05, **p<0.01, ***p<0.001

To explore the functional involvement of these ICD-associated cytokines in regulating endogenous thymus regeneration, we assessed thymus regeneration after SL-TBI in mice with germline deletion of IL-1β signaling (*Il1r1^-/-^*) or IL-18 signaling (*Il18^-/-^* and *Il18r1^-/-^*). Although *Il1r1^-/-^* mice showed no significant changes in their capacity for thymic recovery, suggesting little role for IL-1 signaling, mice deficient for either IL-18 itself or its primary receptor IL-18R1 showed significantly improved thymus regeneration relative to WT controls **(Fig 1f)**. Notably, mice lacking the catalytic domain of Cas-1 also showed significantly increased thymus regeneration **(Fig 1g)**, although to a lesser degree than IL-18-signaling deficient mice possibly reflective of the loss in pro-regenerative effects of DAMP release^15, 16^. Consistent with these findings, administration of recombinant IL-18 (rIL-18) 3 days following SL-TBI – the point at which organ cellularity reaches a nadir and regenerative processes begin to take effect – significantly delays thymic reconstitution **(Fig. 1h)**. Taken together, these data demonstrate that damage-induced activation and release of IL-18 suppresses endogenous thymus repair after acute damage.

We also assessed the therapeutic potential of abrogating IL-18 signaling to improve thymus reconstitution by treating HCT-recipient mice with monoclonal αIL-18 monoclonal antibody. αIL-18 mAb treatment began prior to transplantation and continued for 2-3 weeks following HCT. We hypothesized that capturing not only the acute spike in active IL-18 but also homeostatic IL-18 presence during the initial 2-3 weeks post-transplant would support organ recovery. We find that 50 days post transplantation, mice receiving αIL-18 mAb showed significantly higher thymus cellularity demonstrating that IL-18 can be pharmacologically targeted post transplantation **(Fig. 1i)**. Having established that IL-18 regulates endogenous thymus tissue recovery and that it can be pharmacologically targeted in the context of syngeneic HCT, we set out to identify its source(s) and mechanism of action within the organ.

### IL-18 is produced by discrete populations of hematopoietic and non-hematopoietic stromal cells

Unlike *Il-1*β which is upregulated following inflammasome stimulation, *Il18* is constitutively expressed and is therefore present within the cytoplasm in its pro-form in several cell types awaiting activation by proteolytic cleavage ^18, 28^. To identify the source of IL-18 following acute damage, we investigated *Il18* gene expression from previously published gene expression datasets to narrow the possible source(s) **(Fig S2a)**^29^. From this, we gleaned that, at baseline, *Il18* was not expressed by the large majority of developing thymocytes, and notably, was only expressed within the diverse CD4^-^CD8^-^CD44^+^CD25^-^ DN1 population that includes early T cell precursors, but also myeloid cells, B cells and Innate Lymphoid Cells (ILCs) ^30^. With the knowledge that *Il18* expression was not found in mature thymocytes, we utilized scRNASeq performed on thymus stromal populations that interrogates all non-thymocyte populations by using *Rag2gfp* mice to exclude all Rag2-GFP^+^T cell lineage committed cells **(Fig. 2a, S2b)** ^31, 32^. Utilizing this comprehensive gene expression dataset, we isolated *Il18* expression to non-hematopoietic mesothelial cells and capsular fibroblasts, as well as a subset of dendritic cells (cDC1) and, to a lesser degree, macrophages **(Fig. 2b)**. We could also detect increased cleavage of Cas1 within cDC1 and non-epithelial CD45^-^ cells early after TBI (**Fig. 2c**). This indicates that cDC1 and rare CD45^-^ capsular mesothelial (MEC)/fibroblast (FB) populations meet the qualifications of 1) expressing *Il18* at baseline and therefore are likely to have cytoplasmic pools of inactive pro-IL-18 prior to damage, and 2) increase Cas-1 cleavage following injury necessary for proteolytic cleavage of immature pro IL-18. To functionally interrogate these sources, we generated mice with a specific deletion of *Il18* in conventional dendritic cells using the Zbtb-cre line (*Il18*^Δ^*^cDC^*)^33^. While WT mice increased their levels of IL-18 at day 1 after TBI, *Il18*^Δ^*^cDC^* mice failed to significantly increase levels of activated IL-18 (**Fig. 2d**). To assess the potential contribution of non-hematopoietic stromal cells like MECs and FBs, we performed bone marrow chimeras using WT or *Il18^-/-^* mice as recipients of WT bone marrow. Recipient mice were allowed to recover for 10 weeks following transplantation at which point they received a sublethal dose of TBI and IL-18 levels measured at day 1. Similarly to mice deficient for IL-18 in DCs, these chimera mice demonstrated an increase in IL-18 in WT recipient mice but failed to increase activated IL-18 in *Il18^-/-^* recipients (**Fig. 2e**). However, while not significant, in both of these models we did observe a modest increase in activated IL-18 following injury, consistent with active IL-18 being derived from multiple sources following acute damage.

**Figure 2:**
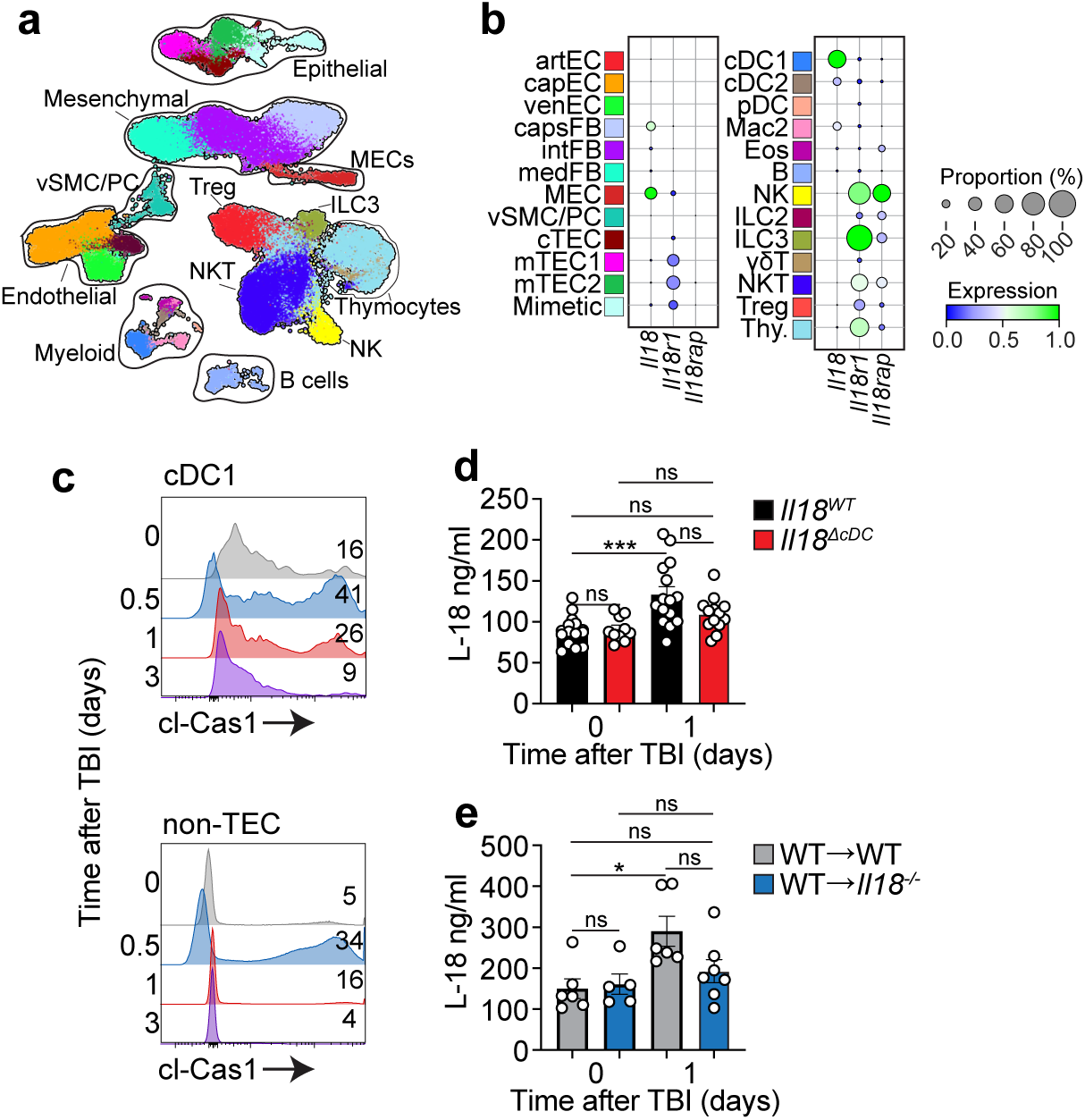
Dendritic cells and non-hematopoietic stroma are sources of damage induced IL-18. **a-b,** Single cell RNA sequencing was performed on (1) non-thymocyte CD45^+^ stromal cells (CD45^+^ *Rag2*-GFP^-^ isolated from female 1-2 mo *Rag2-gfp* mice) and (2) CD45^-^ stromal cells isolated from female 1-2 mo C57BL/6 mice at baseline, 1, 4 and 7 days following SL-TBI. Data was previously integrated together and published in Lemarquis *et al.* (2024). **a**, Integrated UMAP of both datasets showing major clusters and annotation. **b,** Expression of *Il18, Il18r1* and *Il18rap* at baseline. Population abbreviations: arterial Endothelial Cell (artEC), capillary Endothelial Cell (capEC), veinous Endothelial Cells (venEC), capsular Fibroblasts (capsFBs), intermediate Fibroblasts (intFBs), medullary Fibroblasts (mFBs), Mesothelial Cells (MECs), vascular smooth muscle/Pericyte (VSM/PC), cortical Thymic Epithelial Cell (cTEC), medullary Thymic Epithelial Cells (mTEC), proliferating mTEC (mTEC^prol^), cDC (classical Dendritic Cell), Macrophage (Mac), Eosinophils (Eos), Natural Killer (NK), Innate Lymphoid Cells (ILC), Natural Killer T Cell (NKT), Regulatory T cell (Treg), γδ T cell (γδ), Thymocytes (Thy). **c,** cl-Cas1 expression measured by fluorescently conjugated FAM-YVAD-FMK in CD45^+^CD11c^+^MHCII^+^XCR1^+^ cDC1 and CD45^-^EpCAM^-^ non-epithelial stromal cells at days 0, 0.5, 1 and 3 after SL-TBI (n=3-5/group). **d,** Amount of active IL-18 measured by ELISA in female 1-2 mo *Il18^fl/fl^ Zbtb-Cre^-^* (*Il18^WT^*) and *Il18^fl/fl^ Zbtb-Cre^+^* (*Il18*^Δ^*^DC^*) mice at day 0 or 1 following SL-TBI (n=10-16/group). **e,** Female 1-2 mo C57/BL6 WT (WT→WT) or *Il18*^-/-^ (WT→ *Il18*^-/-^) mice were lethally irradiated (2 x 550cGy) and transplanted with 5×10^6^ CD45.1^+^ WT bone marrow hematopoietic cells. 10 weeks after transplantation recipient mice were given a second dose of TBI (sublethal, 550cGy) and active IL-18 was measured on day 1 after this subsequent damage (n=5-7/group). Graphs represent mean ± SEM. *p<0.05, **p<0.01, ***p<0.001

### IL-18 suppression of thymus regeneration is not mediated via direct effect on TECs or hematopoiesis

To determine potential cellular targets for IL-18, we first interrogated previously described transcriptome datasets for expression of the two IL-18R subunit encoding genes^29, 31,32^. We found at baseline that the *Il18r1* subunit was expressed by multiple populations including cortical and medullary thymic epithelial cells (TECs), Tregs, ILCs, NK, and NKT cells (**Fig. 2b, S2a**). TEC expression was especially notable given their role as master regulators of thymus function and regeneration ^34^, although we observed only minimal IL-18R protein expression on TEC subsets **(Fig. 3a-b, S3)** and deletion of *Il18r1* in TECs (using *Foxn1-cre*) demonstrated no change in their capacity for thymus regeneration compared to *Cre^-^*littermate controls **(Fig. 3c)**. Prior work has established that IL-18 can induce hematopoietic stem cell quiescence ^35, 36, 37^. While we do not observe IL18R expression on bone marrow resident precursor populations at baseline **(Fig. 3d-e, S3)**, we did find low levels of IL-18R expression on Early Thymic Progenitors (ETPs), the earliest stage of thymocyte **(Fig. 3a-b, S3)**. Therefore, we explored whether *Il18r1^-/-^* thymocytes showed increased reconstitution capacity relative to WT thymocytes in a competitive transplantation model **(Fig 3f)**. 2 weeks post transplantation, an early timepoint at which initial thymus recovery is measured, CD45.2^+^ *Il18r1^-/-^* cells had no competitive advantage over CD45.2^+^ WT cells in seeding the thymus of CD45.1^+^ WT recipient mice **(Fig 3g)**. Measuring longitudinal contribution of donor derived hematopoiesis by peripheral blood monitoring over 120 days following competitive transplantation showed similar reconstitution in both overall hemopoietic and T cells coming from WT and *Il18r1^-/-^* donor populations **(Fig 3h)**. 120 days post-competitive transplantation, when all hematopoietic cells are of donor origin, we gave recipient mice a subsequent insult of sublethal total body irradiation to directly observe functional regenerative effects of *Il18r1^-/-^* vs. WT thymocytes and saw similar capacities to recolonize the thymus one week later **(Fig. 3i)**. Additionally, these chimeric recipient thymuses showed similar degrees of regeneration by total organ cellularity one week post subsequent injury **(Fig 3j)**.

**Figure 3:**
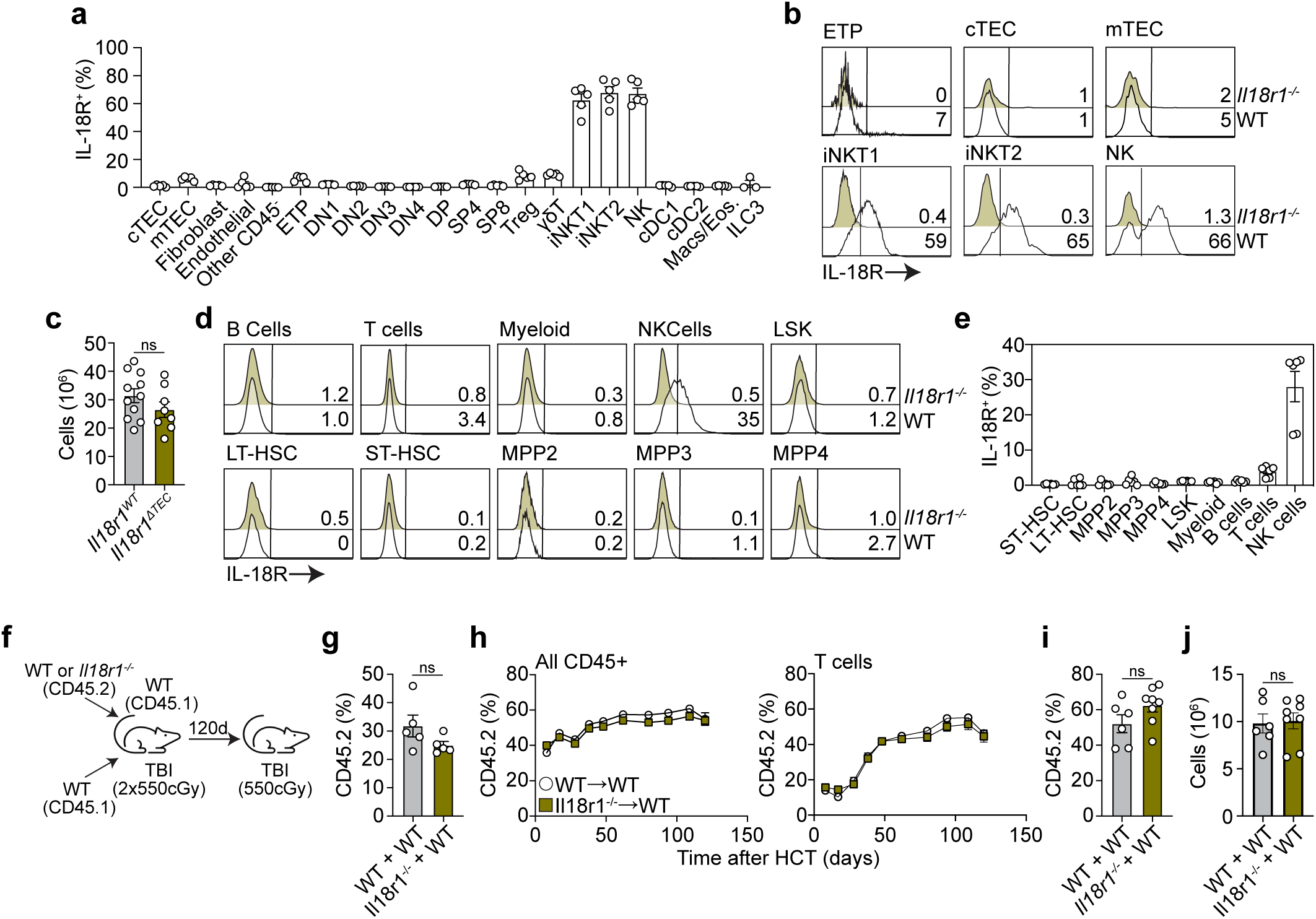
IL-18 suppression of thymus function is not mediated directly through TECs or hematopoietic progenitors. **a-b,** IL-18R expression on thymic cell populations assessed by flow cytometry at baseline (n=3- 5/group). **b,** Concatenated flow plots showing expression of IL-18R on early T-lineage progenitors (ETPs), cortical TECs (cTECs), medullary TECs (mTECs), invariant natural killer cells 1 and 2 (iNKT1/2), and natural killer (NK) cells. Gates were based on expression in *Il18r1^-/-^* mice. **c,** Female 1-2 mo *Il18r1^fl/fl^:FoxN1-Cre^-^*(*Il18r1^WT^*) and *Il18r1^fl/fl^:FoxN1-Cre^+^*(*Il18r1*^Δ^*^TEC^*) were given SL-TBI and thymus cellularity assessed 7 days later (n=8-11/group). **d-e,** Bone marrow populations were measured for IL-18R expression by flow cytometry (n=5-6/group). **f-j, (F)** Female 1-2 mo WT CD45.1^+^ mice were lethally irradiated and transplanted (i.v.) with 2.5×10^6^ WT CD45.1^+^ bone marrow cells and 2.5×10^6^ bone marrow cells from either CD45.2^+^ WT or *Il18r1^-/-^*mice. **g,** Contribution of CD45.2^+^ cells in the thymus at 2 weeks following transplant (n=5/group). **h,** Contribution of CD45.2^+^ cells to peripheral blood total CD45 (left) or T cell (right) reconstitution over 17 weeks post transplantation (n=6-8/group). **i-j,** 17 weeks after transplantation, recipient mice were given a subsequent dose of SL-TBI (550cGy). 7 days later, thymuses were harvested and measured for **(i)** CD45.2^+^ percentage of all thymus CD45^+^ cellularity and **(j)** total thymus cellularity (n=6-8/group). Graphs represent mean ± SEM. *p<0.05, **p<0.01, ***p<0.001

However, despite very little impact *in vivo*, consistent with prior literature, we observed that IL-18 increases thymocyte differentiation and proliferation in studies coculturing *ex vivo* bone marrow derived hemopoietic precursors with the OP9-DLL1 co-culture system **(Fig. S4a-c)** ^38^. From these data we conclude that IL-18 does not suppress thymus function directly via regulating TEC function, nor does it act directly via T-lineage progenitors. Therefore, we hypothesized that IL-18 activity was instead mediated by a thymic intrinsic mechanism.

### Damage Resistant NK Cells Suppress Thymus Regeneration after acute injury

IL-18 can signal through the Il-18R1 subunit but its signaling is potentiated by co-expression of the IL-18 receptor accessory protein Il18rap ^28, 39^. Consistent with their lack of functional signaling after damage, transcriptome datasets indicate that neither TECs nor thymocytes express *Il18rap* (**Fig. 2a, S2a**). Expression of both *Il18r1* and *Il18rap* was largely restricted to ILCs, including NK cells, and NKT cells (**Fig. 2a**). Consistent with this, protein expression suggested NK1.1^+^ populations including iNKT cells and NK cells strongly expressed IL-18R at baseline within the thymus **(Fig 3a-b)**. This was supported by CellChat analysis of scRNASeq which allows for simple and intuitive interactome visualization of sources and targets of IL-18. Targets of IL-18 included NK, NKT and ILC subsets, although NK cell transcriptomes exhibited the strongest aggregate interactome score **(Fig. 4a)**. Following HCT, although both recipient IL-18R^+^ iNKT and NK cells are damage-resistant compared to more abundant thymocyte populations, such that their relative frequency within the organ is significantly higher in the days following transplantation, only NK cells maintained their absolute number with a considerable decrease in NKT cells **(Fig 4b-e)**.

**Figure 4:**
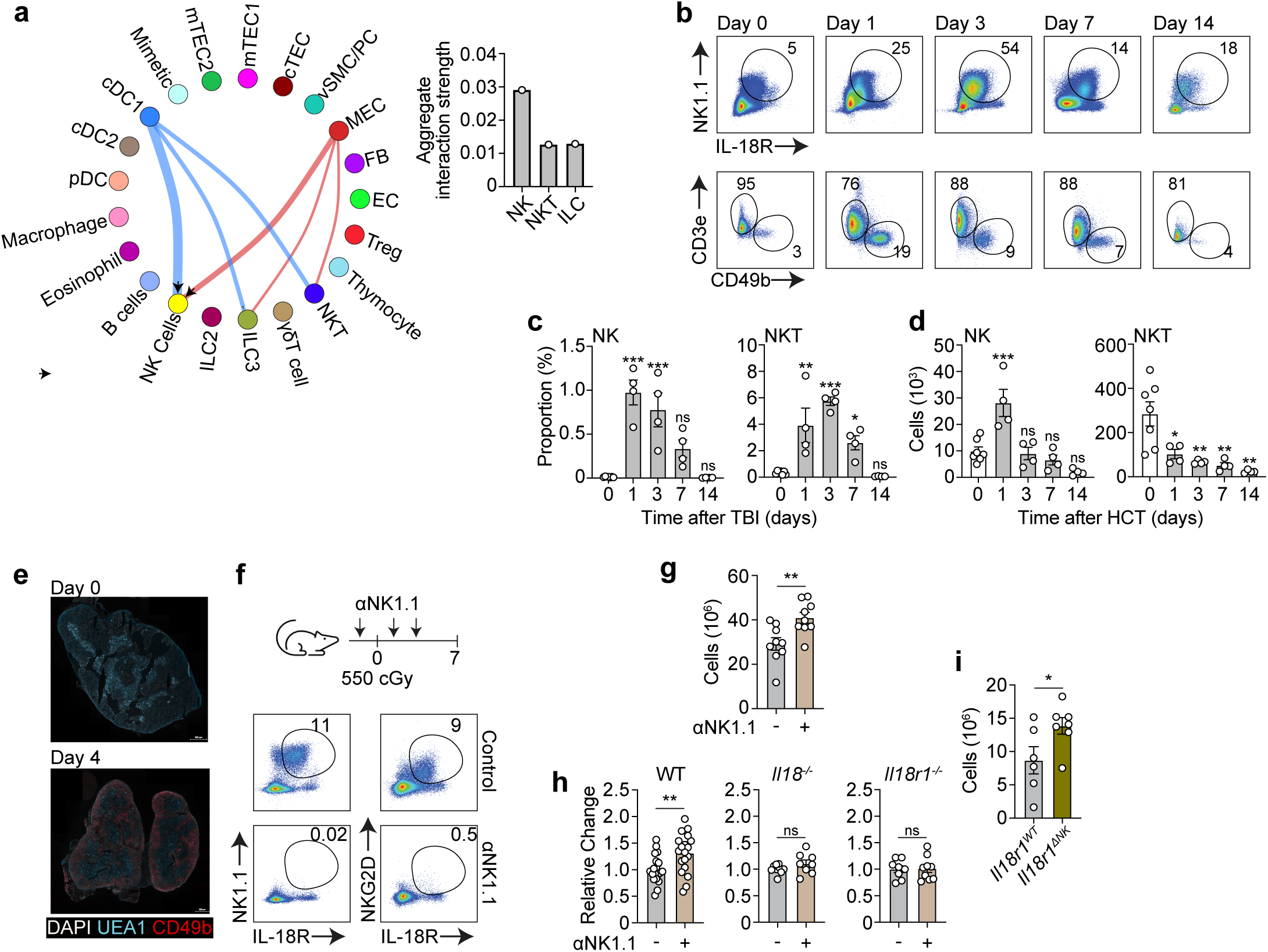
Damage resistant IL-18R^+^ NK cells suppress thymus repair. **a,** CellChat interaction analysis for IL-18 from scRNASeq dataset described in 2A with quantification of the aggregate signal strength coming into any one IL-18 target cell. **b-d,** Female 1-2mo C57BL/6 CD45.2^+^ mice were lethally irradiated and (i.v.) transplanted with 5×10^6^ WT CD45.1^+^ bone marrow cells. **b,** Concatenated flow cytometry plots showing recipient IL- 18R-expressing cells gated on CD45^+^CD45.1^-^CD4^-^CD8^-^(upper) and expression of CD3 or CD49b gated on NK1.1^+^IL-18R^+^ cells at the indicated timepoints after HCT. **c**, Proportion of recipient NK or NKT cells at the indicated timepoints after HCT (n=4-7/group). **d**, Total number of NK or NKT cells at the indicated timepoints after HCT (n=4-7/group). **e,** Female 1-2mo C57BL/6 mice were given SL-TBI and thymus was visualized 4 days later by *in situ* by fluorescent imaging for CD49b^+^ (NK) and UEA-1^+^ (mTECs). **f-g,** Female 1-2 mo C57BL/6 mice were administered 200μg αNK1.1 monoclonal antibody or control PBS (i.p) at days −1, 1 and 3 days post SL-TBI and thymuses assessed at day 7. **f,** Thymus NK1.1^+^IL-18R^+^ (Left) and NKG2D^+^IL-18R^+^ (Right) cells (parent gated on viable CD45^+^CD4^-^CD8^-^ cells) (n=9/group). **g,** Thymus cellularity measured 7 days post SL-TBI (n=9/group). **h,** Female 1-2 mo C57BL/6 WT, *Il18^-/-^*and *Il18r1^-/-^*mice were administered with 200 μg αNK1.1 monoclonal antibody or isotype/ PBS (i.p.) at −1, 1 and 3 days post SL-TBI and thymus assessed at day 7 days. Relative change in thymus cellularity comparing control treated to αNK1.1 treated within each strain (n=8-19/group). **i,** Female 1-2 mo *Il18r1^fl/fl^:Ncr1-Cre^-^* (*Il18r1^WT^*) and *Il18r1^fl/fl^:Ncr1-Cre^+^*(*Il18r1*^Δ^*^NK^*) mice were given SL-TBI and administered with rIL-18 (s.c., 2.5 mg/kg) at day 3. Thymus cellularity measured on day 7 post SL-TBI (n=6-7/group). Graphs represent mean ± SEM. *p<0.05, **p<0.01, ***p<0.001

To assess the role of NK1.1^+^ cells in limiting thymic recovery, we dosed mice with αNK1.1 monoclonal antibody and induced thymus damage by SL-TBI, which showed near complete ablation of thymic IL-18R^+^NK1.1^+^ cells **(Fig 4f).** Importantly, mice depleted of NK1.1^+^ cells had increased thymus cellularity compared to control mice **(Fig. 4g)** suggesting that either iNKT and NK cells are likely mediators of IL-18. Notably, improved regeneration observed upon αNK1.1 antibody treatment in WT thymuses was not recapitulated when performed in either *Il18^-/-^*and *Il18r1^-/-^*mice, suggesting that NK1.1^+^ cell control of thymus recovery is IL-18 dependent **(Fig. 4h)**. To distinguish between the effects on IL-18R^+^ NK vs iNKT cells in regulating thymus regeneration, *Cd1d^-/-^* mice lacking the antigen presenting machinery for iNKT cell development were given SL-TBI to test the effects of thymus suppression in the absence of iNKT cells ^40^. *Cd1d^-/-^* mice had no difference in their thymus cellularity one-week post injury compared to WT control mice **(Fig. S4d**) indicating that IL-18R^+^ iNKT cells, while more abundant than IL-18R^+^ NK cells, do not mediate the IL-18-dependent mechanism limiting thymus regeneration. Finally, to specifically identify NK cells as the target of IL-18 after acute injury, we generated mice with a specific deletion for *Il18r1* in NK cells by using the Ncr1-cre strain, which restricts deletion to cells expressing Nkp46 ^41^. We found that these *Il18r1*^Δ^*^NK^* mice exhibited a significant increase in thymus regeneration after SL-TBI and subsequent rIL-18 supplementation as in Fig. 1h (**Fig. 4i**), strongly indicating that NK cells are mediating the IL-18 response.

### Acute thymic damage activates NK Cells and induces a cytotoxic response

Given that our data demonstrated that IL-18R^+^ NK cells are the main targets of IL-18 following acute thymic damage, we sought to further characterize the functional role of these innate lymphoid cells in regulating organ repair. Analysis of our scRNAseq dataset revealed upregulation following cytoreductive conditioning of a broad program of molecules involved in NK cell function, including the effectors *Ifng*, *Prf1, Gzma* and *Gzmb* **(Fig 5a)**. Comparison with other potential IL-18R^+^ targets such as NKT cells revealed no such program (**Fig. 5a**). Protein analysis of thymus NK cells 3 days following HCT-conditioning supported the transcriptome with increased NK expression of IFNγ, Granzyme B, Perforin, and surface marker NKG2D demonstrating broad induction of NK cell activation and cytotoxicity **(Figs 5b)**. Notably, thymus IL-18R^+^ NK cells make up the largest population of Perforin^+^ cells within the thymus following damage **(Fig 5c)**. Consistent with this, we also found a significant increase in the global thymic levels of IFNγ, Granzyme B, and Perforin early after HCT-conditioning (**Fig. 5d**).

**Figure 5:**
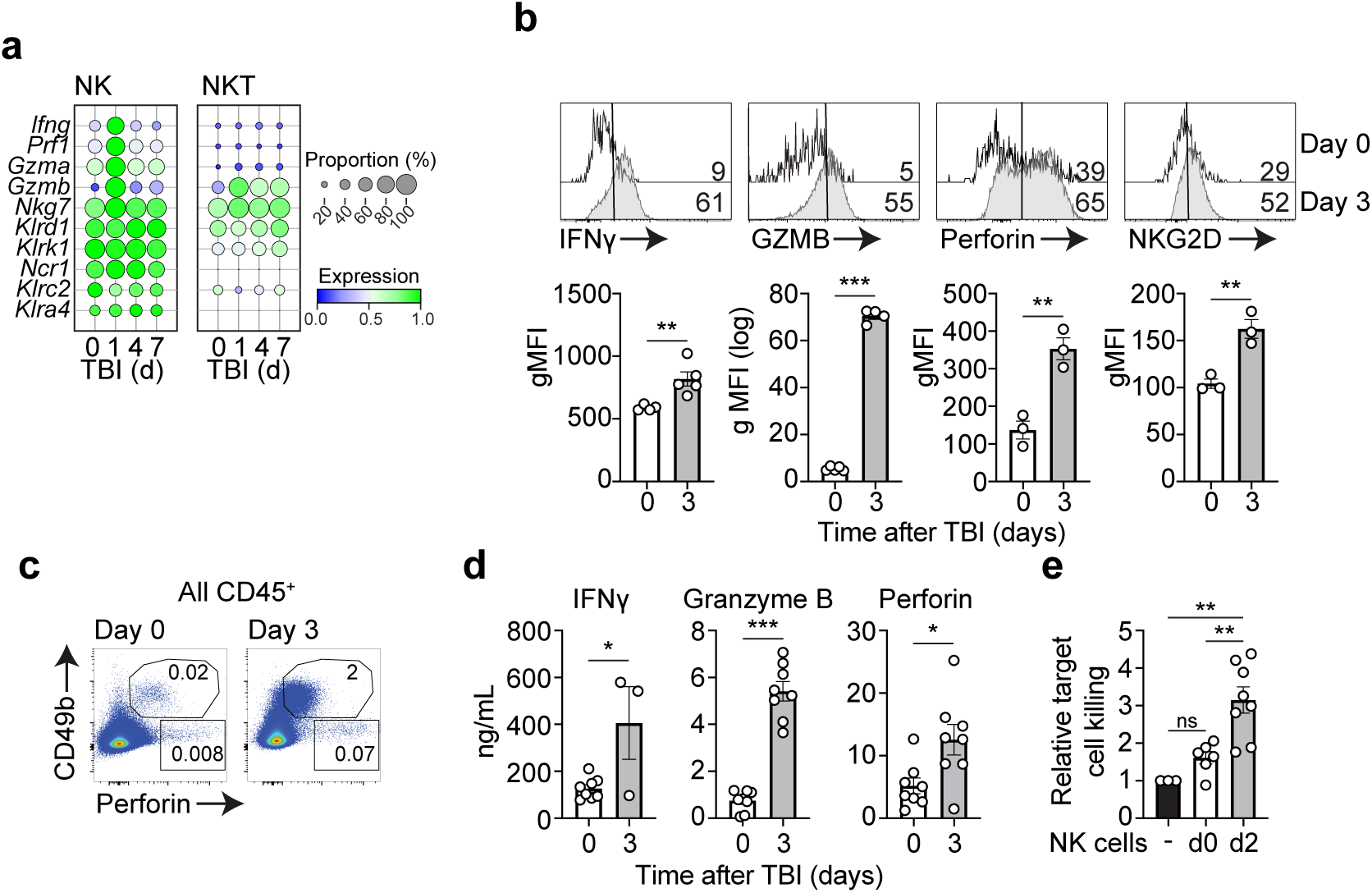
HCT conditioning activates thymic NK cells. **a,** NK and iNKT cell gene expression of cytotoxicity factors *Ifng, Prf1, Gzma, Gzmb* and activation markers *Nkg7, Klrg1, Klrk1* at days 0, 1, 4 and 7 post SL-TBI taken from scRNASeq dataset described in 2A. **b,** IL-18R^+^NK1.1^+^CD49b^+^ NK cell expression of *Ifng*-GFP, Granzyme B (GzmB), Perforin and surface NKG2D at days 0 and 3 following SL-TBI in female 1-2 mo C57BL/6 WT or *Ifng*-reporter (GREAT) mice (n=3-5/group). **c,** Concatenated plots showing perforin and IL-18R expression in CD45^+^ cells at baseline or 3 days post SL-TBI. **d,** Amount of thymic IFNγ, Granzyme B, and Perforin measured by ELISA from female 1-2 mo C57BL/6 mice at day 0 and 3 post SL-TBI (n=3-8/group). **e,** NK1.1^+^IL-18R^+^TCRβ^-^CD49b^+^ NK cells were FACS purified from female 1-2 mo C57BL/6 mice at baseline (day 0) or 2 days post SL-TBI and cocultured with cell-trace labeled RMA-S target cells at a 2:1 effector to target ratio. RMA-S Target cell Annexin V expression was measured at 5 hours post co-culture and cell death assessed relative to RMA-S cells cultured without NK cells present (n=3-8/group). Graphs represent mean ± SEM. *p<0.05, **p<0.01, ***p<0.001

Having demonstrated that NK cells are stimulated by cytoreductive conditioning induced thymic damage, we attempted to determine whether this increased NK activation profile directly resulted in increased NK cytotoxicity and killing of target cells. To address this aim, we cocultured IL-18R^+^ NK cells FACS purified from either undamaged thymuses or from acutely involuted thymuses 2 days following HCT conditioning with RMA-S target cells. These *ex vivo* cytotoxicity assays revealed that NK cells from HCT-conditioning damaged thymuses were significantly more cytotoxic than those from the undamaged thymus **(Fig 5e)**. Taken together our data show that acute damage to the thymus such as that caused by HCT-conditioning activates resident NK cells, increasing their cytotoxicity and capacity for killing nearby target cells.

### IL-18 mediates the NK cell effector program after acute damage

Although the role for IL-18 was originally largely restricted to its induction of IFNγ – in fact, IL-18 was first known as IFNγ inducing factor ^42^ – recent evidence suggests a broader role in activating cells such as NK cells ^17, 28^ Consistent with this broader role we observed that following cytoreductive conditioning, expression of the activation and effector genes *Klrk1, Nkg7, Gzma, Gzmb, Pfr1* and *Ifng* in NK cells correlated with the expression of the restricted coreceptor *Il18rap* shortly following acute damage, which was not the case in NKT cells **(Fig 6a)**. This data strongly suggests that IL-18 signaling mediates the activation of this program. However, to functionally determine whether IL-18 directly induced activation of thymic NKs, we administered rIL-18 to mice at baseline that have not been exposed to any damaging stimuli and assessed NK cell derived cytotoxic factors 2 days later. Although in this model thymic size was unaffected 2 days following administration, rIL-18 alone did increase the number of NKs within the thymus and their expression of the effector molecules Perforin and IFNγ **(Figs. 6b-d)**. Furthermore, NK cells isolated from thymuses of mice receiving rIL-18 3 days post cytoreductive conditioning and harvested 4 days thereafter (as in Fig 1H) showed significantly higher cytotoxicity than thymic NK cells from mice receiving control PBS **(Fig. 6e)**. Together, these findings suggest that IL-18 release following HCT-conditioning increases thymus NK cell cytotoxicity.

**Figure 6:**
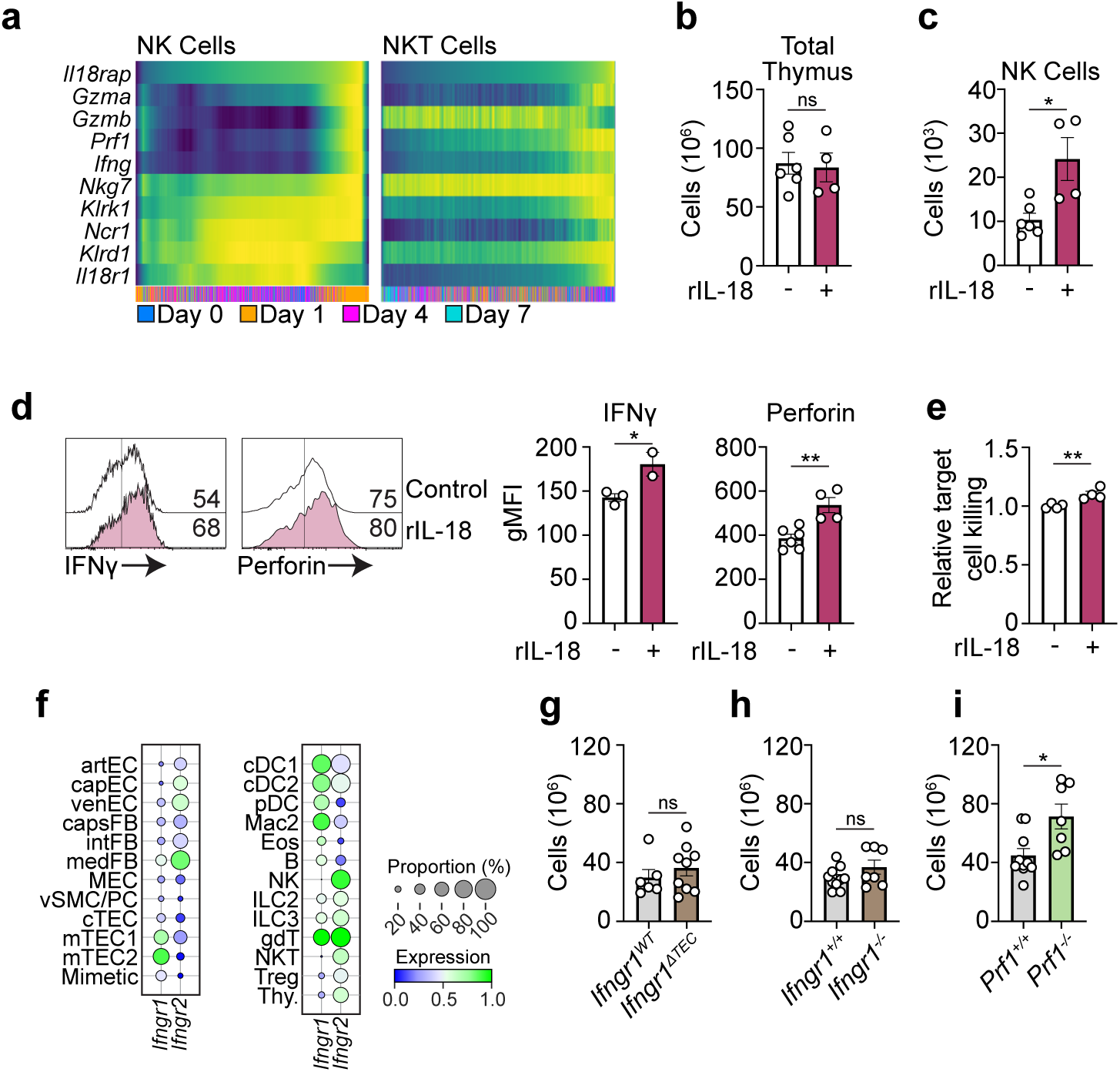
IL-18 stimulation of cytotoxic NK cells suppresses thymus regeneration. **a,** Heat map of thymus NK and iNKT cell gene expression of *Gzma, Gzmb, Prf1, Ifng, Nkg7, Klrk1, Ncr1, Klrd1 and Il18r1* 1, 4 and 7 days following SL-TBI relative to expression of *Il18rap* (taken from scRNASeq dataset described in 2A). **b-d,** Female 1-2 mo C57BL/6 WT or *Ifng*- reporter mice were given rIL-18 (s.c., 2.5mg/kg) or PBS and total thymus cellularity (**b**) or IL- 18R^+^ NK cell number (**c**) assessed on day 3 (n=4-6/group). **d**, Expression of *Ifng*-GFP or Perforin at day 3 (n=2-6/group). **e,** Female 1-2mo C57BL/6 mice were given SL-TBI and 3 days later administered with rIL-18 (s.c., 2.5mg/kg) or PBS. ON day 7 NK1.1^+^IL-18R^+^TCRβ^-^CD49b^+^ NK cells were then FACS purified and cocultured with cell-dye labeled RMA-S target cells at a 5:1 Effector to Target ratio. RMA-S target cell Annexin V expression was measured 5 hours post co-culture (n=4/group). **f,** *Ifngr1* and *Ifngr2* expression at baseline taken from the scRNASeq dataset described in 2A. **g,** Female 1-2 mo C57BL/6 *Ifngr^fl/^fl:Foxn1-Cre^-^*(*Ifngr^WT^*) and *Ifngr^fl/^fl:Foxn1-Cre^+^* (*Ifngr*^Δ^*^TEC^*) mice were given SL-TBI and thymus cellularity measured on day 7 (n=6-9/group). **h,** Female 1-2 mo C57BL/6 WT (*Ifngr ^+/+^*) or *Ifngr1^-/-^* mice were given SL-TBI and thymus cellularity was measured at day 7 (n=7-8/group). **I,** Female 1-2 mo C57BL/6 WT (*Prf ^+/+^*) and *Prf ^-/-^* mice were given SL-TBI and thymus cellularity was measured at day 7 (n=7- 10/group). Graphs represent mean ± SEM. *p<0.05, **p<0.01, ***p<0.001

The most widely studied function of IL-18 is its capacity to induce IFNγ production from T cells and NK cells ^43^ Our scRNAseq dataset suggested widespread expression of IFNγ receptor genes, including on crucial stromal subsets of thymic epithelial cells (**Fig. 6f**); a pathway that has previously been implicated in mediating TEC cell death in acute graft versus host disease after HCT ^44^ This led us to hypothesize that IL-18 activated NK cell released IFNγ is responsible for suppressing thymus regeneration via direct targeting of TECs. To address this, we generated mice with a TEC-specific deletion for *Ifngr1;* however, absence of IFNγR on TECs identified no difference in thymic regenerative capacity, suggesting that TECs are not a direct target of IFNγ after injury (**Fig. 6g**). To assess if IFNγ could be affecting other cells, we assessed regeneration after SL-TBI in mice with a germline deletion for *Ifngr*, which similarly did not show significantly higher thymus cellularity one week post conditioning compared to WT controls **(Fig 6h)**. NK cell mediated killing involves degranulation releasing preformed cytotoxic proteins, mainly granzymes and perforin ^45^ Mice lacking Perforin (*Prf^-/-^*) did in fact show significantly improved thymus regeneration compared to wildtype control mice **(Fig. 6i)**. Therefore, we demonstrate that Perforin-dependent direct cytotoxicity of NK cells is capable of suppressing thymus repair post-injury.

### Cytotoxic NK Cells Aberrantly Target Thymic Epithelial Cells

Ly49 family receptors on NK cells recognize stochastically expressed MHC-I on cells and its binding inhibits NK cytotoxicity ^46^. Several viruses evolved to express proteins interfering with MHC-I antigen presentation, and NK cells are equipped to recognize virally infected cells’ downregulation of MHC-I to mediate their killing. Similarly, downregulation of MHC-I on tumor cells evading CD8^+^ T cell mediated immune surveillance are also targeted by NK cells ^46^. Having established that NK cell cytotoxicity suppresses thymus regeneration, we sought to identify populations that may be targeted by NK cells following cytoreductive conditioning. We observed significant downregulation of MHC-I component H2-kb in both cTECs and mTECs following acute thymus damage **(Fig 7a)**. TECs not only downregulate NK cytotoxicity inhibitory factor MHC class 1, but they also upregulate NK stimulating NKG2D ligand RAE-1 following cytoreductive conditioning **(Figs 7b)**. Together, downregulation of H2-kb and upregulation of NKG2D ligands makes TECs vulnerable to NK cell cytotoxicity. Given the crucial function of TECs during normal T cell development as well as thymic regeneration^34^, we hypothesized that TECs were targeted by IL-18 triggered NKs following cytoreductive conditioning, which would ultimately serve to suppress thymus function. Consistent with the differential degree of MHC-I downregulation and Rae1 upregulation, we found that mTEC cellularity was significantly greater in *Il18*^-/-^ mice while cTEC cellularity was not significantly affected 7 days post SL-TBI **(Fig 7c)**. We also found a similar differential effect on TEC subsets in mice whose thymus recovery was improved by treating with αNK1.1 mAb (as in Fig 3I) **(Fig 7d)**. These data suggested that IL-18 triggered cytotoxic NK cells preferentially mediate mTEC cell destruction.

**Figure 7:**
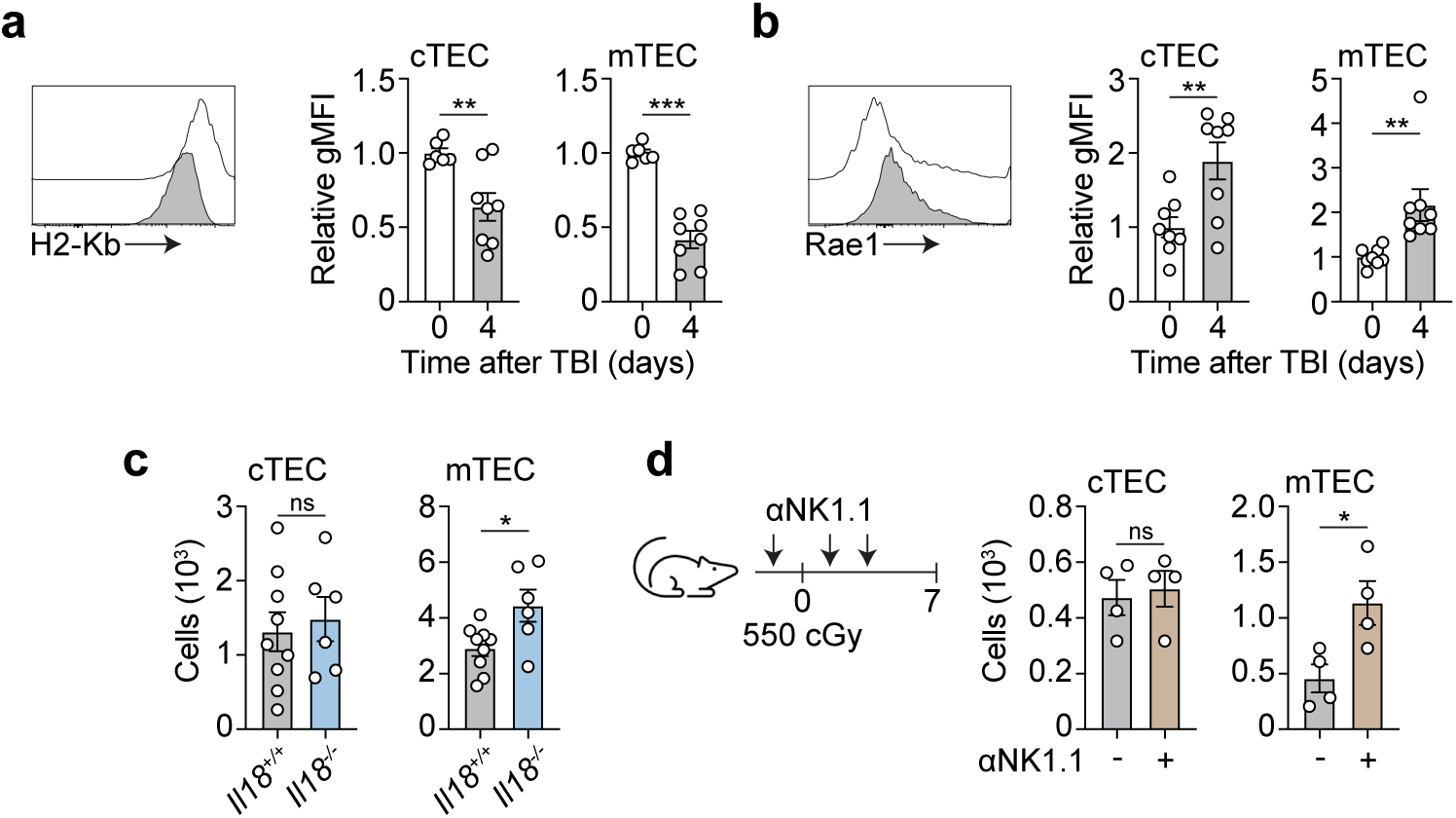
Cytotoxic NK cells aberrantly target Thymic Epithelial Cells. **a-b,** Expression of MHC-I (**a**) and pan-Rae1 (**b**) at day 0 and 4 following SL-TBI of 2mo C57BL/6 mice. Concatentaed flow plots gated on all TECs (CD45^-^EpCAM^+^MHC-II^+^) and graphs showing MFI within specific cTEC and mTEC populations (n=6-8/group). **c,** Female 1-2 mo C57BL/6 WT and *Il18 ^-/-^*mice were given SL-TBI and thymus assessed at day 7 for number of CD45^-^EpCAM^+^MHC-II^+^Ly51^+^UEA-1^-^ cTEC and CD45^-^EpCAM^+^MHC-II^+^Ly51^-^UEA-1^+^ mTEC were quantified (n=6-9/group). **d,** Female 1-2 mo C57BL/6 mice were administered 200μg αNK1.1 monoclonal antibody or PBS at −1, 1 and 3 days post SL-TBI. Thymus was analyzed on day 7 days for number of cTEC and mTEC (n=4/group). Graphs represent mean ± SEM. *p<0.05, **p<0.01, ***p<0.001

Taken together, we have identified that following cytoreductive conditioning, increased cl-Cas-1 leads to the release of activated IL-18 which triggers the cytotoxicity of organ-resident NK cells, and that these NK cells target TECs, particularly mTECs, to suppress thymus recovery and T cell reconstitution. Furthermore, these data also demonstrate that IL-18 abrogation holds promise as a therapeutically feasible strategy that can improve thymus recovery post-HCT.

## DISCUSSION

In this study, we have identified a crucial role for IL-18 in limiting thymic regeneration following acute damage via stimulation of tissue-resident NK cells, which aberrantly targeted key stromal populations such as mTECs. Despite its importance for generating and maintaining a pool of naïve T cells, the thymus is extremely sensitive to multiple forms of acute injury including everyday insults like stress and infection, but also more profound damage like cytoreductive chemotherapy and ionizing radiation. We recently reported that thymocyte depletion following ionizing radiation is mediated by sharp rises in both caspase-1-mediated immunogenic cell death (ICD) as well as caspase-3 mediated apoptosis in thymocytes^16^. Products of this lytic cell death including Zinc and ATP support thymic regeneration post-injury by directly or indirectly acting on TECs, the primary supporting stromal cell supporting T cell development^15, 16, 47^. Here, we demonstrate that increased ICD not only occurs following radiation injury, but also during the depletion phase of other acute thymus insults including common chemotherapy regimens, corticosteroid treatment and sepsis.

During infection, myeloid cells such as DCs and macrophages release caspase-1 activated IL-18 that stimulates potent IFNγ production by CD4^+^ Th1, CD8^+^ and NK cells^48^. As an acute-phase effector response, constitutive expression in cells lead to cytoplasmic stores of the inactive immature form of IL-18 that await proteolytic cleavage^28, 49^. Our data confirms that in the thymus, hematopoietic constitutive expression of *Il18* was restricted largely to cDC1s. Indeed, abrogation of *Il18* expression specifically within DCs did mitigate the rise in IL-18 following conditioning. Interestingly, we could also detect a relatively high baseline amount of cl-Cas-1 in cDC1 and low levels of active IL-18 in the organ, which could be consistent with a previously reported homeostatic role for IL-18 in maintaining a population of thymus-resident regulatory T cells^50^. However, protein still rose in cDC conditional *Il18* knockout mice consistent with there being alternate sources of activated IL-18 post-injury suggesting other sources. Supporting the possibility of an alternate source of IL-18, we also found constitutive expression of IL-18 in non-hematopoietic cells such as capsular fibroblasts and mesothelial cells lining the organ parenchyma. This expression was supported by increased cl-Cas1 in the non-hematopoietic compartment after TBI and functionally confirmed by bone marrow chimeras that showed decreased levels of protein following damage in *Il18^-/-^* recipients. This alternate source of IL-18 is consistent with reports suggesting nonhematopoietic sources are functional contributors of intestinal integrity regulating IL-18^51, 52^. This study demonstrating that capase-1 dependent IL-18 is a novel suppressant of thymus regeneration is also supported by previous work in which mice deficient for NLRP3 inflammasome upstream of caspase-1 showed improved thymus function^53^.

IL-18 has been shown to regulate intestinal barrier function via epithelial cell maturation and function^51, 52, 54^. Given existing parallels between intestinal epithelium and TECs, we first explored the possibility that IL-18 signaling via TECs may directly inhibit their function. However, although there was detectable *Il18r1* expression within various mTEC subsets, neither the *Il18rap* coreceptor, nor significant levels of IL-18R protein expression was found on any TEC subset. Crucially, conditional genetic deletion of *Il18r1* within *Foxn1* expressing TECs did not alter thymus recovery following TBI. One additional mechanism that has been demonstrated for IL-18 is in the regulation of bone marrow hematopoiesis^36, 37^ which, given the fundamental requirement for bone marrow progenitor seeding of T cell development, was a distinct possibility for a mechanism underlying the role of IL-18 in thymic regeneration. However, we could detect no difference in the capacity for *Il18r1* deficient hematopoietic stem and progenitor cells to 1) reseed the recovering thymus shortly following transplantation and 2) reconstitute overall hematopoietic and T cell populations longitudinally 120 days post-transplant. Furthermore, recipients receiving WT and *Il18r1^-/-^* donor cells showed similar capacity for recovery after a subsequent insult of SL-TBI 4 months following transplantation. Based on these findings, and despite prior evidence for an effect of IL-18 on epithelial cells in the gut and on hematopoietic progenitors in the BM, these mechanisms did not explain the IL-18 suppression of tissue repair in the thymus.

IL-18 canonically signals through T cells, NKT and NK cells to mediate a TH1 response, primarily by inducing expression of IFNγ^18, 28^. Indeed, although there was some expression of *Il18r1* on TECs, ETPs, Tregs, and ILCs, by far the greatest expression was restricted to NKT cell progenitors and NK cells. While we found that depletion of both NK and NKT populations with αNK1.1 mAb improved thymus function after damage. In contrast with studies implicating NKT cells in regeneration^11^, in our hands mice deficient for NKT cells did not show any alteration in thymic repair. Furthermore, expression of the coreceptor *Il18rap*, which significantly potentiates IL-18 signaling^28, 39^, was largely absent on NKT cells and was restricted in the thymus to NK cells. Importantly, specific deletion of *Il18r1* in NK cells revealed improved thymus regeneration. However, despite previous studies linking IFNγ with TEC cell death during graft versus host disease^44^, surprisingly IFNγ did not seem to be the effector molecule used by NK cells to limit thymic regeneration. Instead, IL-18 induced a broad activation program including upregulation of effectors of direct cytotoxicity such as granzymes and perforin, and mice deficient for perforin demonstrated improved thymus reconstitution. IL-18 requires co-stimulation with either IL-12 or IL-15 to activate of CD4^+^ Th1 and CD8^+^ T cytotoxic cells, however, recent reports suggest that IL-18 alone is capable of stimulating a broad activation program in NK cells^17^, and accordingly we found that sole rIL-18 administration increased NK cell Perforin, GzmB and IFNγ production.

This study represents the first recognition of tissue-resident NK cells negatively regulating thymus function. In fact, donor NK cells are reportedly beneficial in the setting of HCT, promoting engraftment, reducing GVHD by targeting HLA-mismatched antigen presenting cells, and increasing TEC proliferation^55, 56^. However, we suggest a different mechanism whereby radio-resistant *recipient* NK cells are activated as a consequence of damage and counteract thymus regeneration. While ionizing radiation has been shown to induce expression of MHC class I in tissues such as gut^57, 58^, this was not the case in TECs which showed both a decrease in MHC I expression but also upregulation of the NK-activating receptor NKG2D ligand Rae1, making them vulnerable to cytotoxicity in an HLA-mismatch independent fashion. However, the skewing toward mTEC targeting could be explained by their higher rate of cell turnover and proliferation, contributing to their sensitivity to radiation damage, both of which can be drivers for NKG2D ligand expression^59, 60, 61, 62^. Therefore, there is a distinction between the pro and anti-reparative roles of functions of donor and recipient NK cells, respectively. Furthermore, our work is consistent with reports showing that IL-18 can stimulate NK cells that aberrantly target epithelium during HSV-2 infection and delay re-epithelization in wound healing, as well as with NK cell directed cytotoxicity targeting HSPCs that upregulate NKG2D ligands in Fanconi Anemia due to genotoxic stress^19, 20, 21, 60^. Our study therefore contributes to a growing body of literature revealing a role for NK cells in regulation of tissue injury and recuperation.

This work positions IL-18 as a potential therapeutic target for improving thymus function after acute injury such as HCT conditioning. However, given its reported context-dependent effects of IL-18 in GVHD, along with its emerging promise in immunotherapy, careful examination of its abrogation will be required to balance its pro-reparative and graft-versus-tumor effects^18, 63, 64, 65, 66, 67, 68^. In summary, this study identifies a novel pathway regulating T cell immune reconstitution following acute thymus damage and presents multiple opportunities for potential therapeutic targeting to improve T cell reconstitution not only in HCT patients, but also those exposed to other acute thymus damages due to chemotherapy, stress, and infection.

## METHODS

### Mice

Inbred male and female C57Bl/6J (000664) and B6 CD45.1 (002014) mice were obtained from Jackson Laboratories (Bar Harbor, USA). *Il1r1^-/-^*(003245), *Il18^-/-^*(004130), *Il18r1^-/-^*(004131), *Casp1*Δ*10* (032662), *CD1d^-/-^* (008881), *Ifngr^-/-^* (003288), *Prf1^-/-^*(002407), *Rag2-eGFP* (005688) and *GREAT* (“interferon-<u>g</u>amma reporter with <u>e</u>ndogenous poly<u>A t</u>ranscript”) mice were obtained from Jackson Laboratories and bred in house. Il18 flox mice (*Il18^fl/fl^*) were obtained from R. Nowarski (Harvard Medical School) and R. Flavell (Yale School of Medicine) and crossed in house to *Zbtb46 Cre^+^* obtained from Jackson Laboratories (032662) to generate *Il18^fl/fl^ Zbtb Cre^+^*(*Il18*^Δ^*^DC^*) mice. *Il18r1^fl/fl^* mice were obtained from G. Trinchieri (NCI) and crossed to *Foxn1 Cre^+^* mice obtained from Jackson Laboratories (018448) and *Ncr1 Cre^+^* obtained from K. Barry (Fred Hutch Cancer Center) to generate *Il18r1^fl/fl^ Foxn1 Cre^+^* (*Il18r1*^Δ^*^TEC^*) and *Il18r1^fl/fl^ Ncr1 Cre^+^* (*Il18r1*^Δ^*^NK^*), respectively. *Ifngr^fl/fl^* (025394) and *Foxn1 Cre^+^* (018448) mice were obtained from Jackson Laboratories and crossed to generate *Ifngr^fl/fl^ Foxn1 Cre^+^*(*Ifngr*^Δ^*^TEC^*) mice. All experimental mice were between 6-10 weeks old. Mice were maintained at the Fred Hutchinson Cancer Research Center (Seattle, WA), and acclimatized for at least 2 days before experimentation, which was performed per <u>Institutional Animal Care and Use Committee</u> guidelines.

### Cell Isolation

Single cell suspensions of freshly dissected thymuses were obtained and enzymatically digested using 0.15% Collagenase D (Sigma 11088882001) and 0.1% DNase 1 (Sigma 10104159001) in DMEM, as previously described.^8^ Cellularity was calculated using the Z2 Coulter Particle and Size Analyzer (Beckman Coulter, USA). For studies sorting rare populations of cells in the thymus, multiple identically-treated thymuses were pooled so that sufficient numbers of cells could be isolated; however, in these instances, separate pools of cells were established to maintain individual samples as biological replicates. BM was flushed from femors and tibias and then filtered through a 70 μm filter. Peripheral blood was collected into EDTA capillary pipettes (Fisher Scientific). Red Blood Cell Lysis was performed with ACK Lysis Buffer (A1049201, Fisher Scientific).

### Flow Cytometry

Cells were stained with the following antibodies for analysis: CD45 (565967, BD Biosciences), CD31 (102434, Biolegend), CD140a (135907, Biolegend), MHC-II (107620, Biolegend), EpCAM (46-5791-82, BD Biosciences), Ly-51 (740882, BD Biosciences), UEA-1 (ZC0426, Vector Labs), CD8a (100714, Biolegend), CD4 (565709, BD Biosciences), TCR-β (109239), CD3e (100232, Biolegend), CD25 (102030, Biolegend), CD44 (612799, BD Biosciences), NK1.1 (108753, Biolegend), CD49b (561067, BD Biosciences), c-Kit (105811, Biolegend), TCR-γ𝛿 (118107, Biolegend), CD1d PBS-57 Tetramer (NIH tetramer Core), CD11c (35-0114, Tonbo), CD11c (612796, BD Biosciences), CD11b (741722, BD Biosciences), XCR1 (148225, Biolegend), B220 (103232, Biolegend), CD90.2 (105331, Biolegend), CD127 (50-1271, Tonbo), Sca-1 (122527, Biolegend), CD135 (135305, Biolegend), CD150 (46-1502-82, eBioscience), CD48 (103427, Biolegend), NKG2D (562800, BD Biosciences), H2-kb (116525, Biolegend), Pan Rae-1 (FAB17582P, R&D), IL-18R (25-5183-82, Thermo Fisher), IL-18R (25-5183-82, Thermofisher). Following fixation and permeabilization (554714, BD Biosciences), cells were stained with the following antibodies: Perforin (154315, Biolegend), Granzyme B (MHGB04, Thermofisher). Annexin V staining (640920, BioLegend) was performed in Annexin V binding buffer (422201, BioLegend). Flow cytometric analysis was performed on a Symphony S6 (BD Biosciences) and cells were sorted on an Aria II (BD Biosciences) using FACSDiva (BD Biosciences) or FlowJo (Treestar Software).

### In vivo Acute Damage Models

To induce thymus damage, mice were given sublethal total body irradiation at a dose of 550 cGy from a cesium source mouse irradiator (model here) with no hematopoietic rescue. Other models of thymus damage included i.p. injection of 20mg/kg Dexamethasone (Sigma-Aldrich D2915), 200mg/kg Cyclophosphamide (University of Washington Medical Pharmacy) or 1.5mg/kg LPS (Invivogen tlrl-eblps). For *in vivo* studies of rIL-18 administration, C57Bl/6J or GREAT mice were either given (s.c.) 2.5 mg/kg rIL-18 in the absence of other thymus damaging treatment (Day 0) or 3 days post SL-TBI.

### In vivo Depletion and Transplantation Studies

To perform NK1.1^+^ cell depletion studies, mice were injected (i.p.) with 200ug (10mg/kg) αNK1.1 mAb (BioXCell BE0036) at days −1, 1 and 3 post-SL-TBI. B6 HCT recipients received 1100 cGy TBI (2 × 550 cGy) prior to transplantation and within 24hrs received i.v. injection of 5 to 10 × 10^6^ bone marrow (BM) cells. To perform IL-18 abrogation experiments, mice were dosed (i.p.) with 200ug (10mg/kg) αIL-18 mAb (BioXCell BE0237) at days −1, 1, 3, 6, 9, 12, 15 and 18 post- transplantation.

### Protein quantification

For detection of supernatant active IL-1β, active IL-18, Granzyme B, Perforin, IFNγ, and mature Cas-1 (Figs 1B-C, 5C, and S1) were obtained by mechanically digesting thymic in defined volumes of buffer. The resulting supernatant was quantified using cytokine specific ELISA kits (IL-1β Invitrogen #88-7013-22; IL-18 Thermo Fisher #BMS618-3; GzmB R&D #DY1865; Perforin Novus Biologicals # NBP3-00452; IFNγ Thermo Fisher #KMC4021; mature Cas-1 Adipogen #AG-45B-0002-KI01) and absorbance was measured on Tecan Spark 10M (Tecan, Switzerland).

For detection of whole organ active IL-18 and IL-18 Binding Protein (1B only post- cyclophosphamide IL-18, 1D, 2D-E) thymuses were homogenized in RIPA buffer (25mM Tris pH 7.6, 150nM NaCl, 1% NaCl, 1% NP-40, 0.1% SDS, 0.05% sodium deoxycholate, 0.5mM EDTA) with protease inhibitors (Thermo Fisher A32955), using a homogenizer 150 (Fisher Scientific) and normalizing by mass at a concentration of 20 mg thymus tissue/mL RIPA buffer. The resulting lysates was quantified using cytokine specific ELISA kits (IL-18 Thermo Fisher #BMS618-3 and IL-18 BP Abcam # ab254509), and absorbance was measured on the Tecan Spark 10M (Tecan, Switzerland).

### In vitro cell culture

Co-culture experiments were performed by plating 50,000 *ex vivo* FACS purified bone marrow Lineage^-^ selected or Lineage^-^ Sca-1^+^ c-Kit^+^ FACS purified cells onto 6 well-plates confluent with OP9-DLL1^GFP^ cells in OP9 media previously described.^15, 69^ Co-cultures were performed in the presence of 5 ng/mL Flt-3L (Peprotech, 250-31L) and 1 ng/mL IL-7 (Peprotech 217-17) and either 0 ng/mL, 1ng/mL or 10 ng/mL rIL-18 (Biolegend 767008). Equal volumes of non-adherent cells were assessed by flow cytometry for differentiation at 10, 14 or 21 days post co-culture.

### Cytotoxicity Assays

Cytotoxicity assays were performed by co-culturing NK1.1^+^ IL-18R^+^ CD49b^+^ TCRβ^-^ NK cells FACS isolated from either undamaged (Day 0) or 48 hours post SL-TBI with Cell-trace Yellow (Thermofisher C34573) labeled RMA-S cells at a 2:1 Effector to Target ratio in RPMI supplemented with 10ng/mL rIL-15 (Biolegend 566302). Co-cultures were incubated at 37° degrees for 5 hours, after which cell death of Cell Trace Yellow (CTY) labeled RMA-S cells was assessed by flow cytometry according to Annexin V (Biolegend 640920) expression.

### Imaging

Thymuses were mounted in optimal cutting temperature coumpound (OCT, TissueTek) and snap-frozen, and 10um sections were fixed in a 1:1 methanol/acetone solution. Sections were stained with anti-CD49b (Biolegend 108901), UEA-1 (Vector Laboratories), and detected with Alexa 568 (Thermo Fisher Scientific).

4′,6-diamidino-2-phenylindole (DAPI) was stained at a concentration of 300 nM, and sections were mounted with Vectashield Plus (Vecto Laboratories, Burlingame, CA) and imaged using a TissueFAX Plus (Tissuegnostic, Tarzana, CA). Fluorescent images of whole thymic tissue were analyzed using ImageJ 1.5i.

### Single Cell RNA Sequencing

Previously generated and published single cell RNA-seq datasets of thymic CD45- non- hematopoietic cells (GSE240016; 50,890 cells) and RAG2-CD45+ hematopoietic cells (GSE244673; 37,879 cells) from 2-month-old mice at steady state and days 1, 4 and 7 after SL- TBI were used for this study^31, 32^. The CD45- dataset can be viewed at https://thymosight.org/ along with all previously published thymus single cell sequencing datasets.

### Statistics

Statistical analysis between two groups was performed with unpaired two-tailed t test. Statistical comparison between 3 or more groups in Figures was performed using a one-way ANOVA with Dunnett’s multiple comparison test (2D-E, 4C-D). All statistics were calculated using Graphpad Prism and display graphs were generated in Graphpad Prism or R. Information on replicates, error bars and statistical significance can be found in the figures and their corresponding legends.

## ACKNOWLEDGMENTS

We gratefully acknowledge the assistance of the Flow Cytometry and Comparative Medicine Core Facilities; and the support of the Immunotherapy Integrated Research Center at the Fred Hutchinson Cancer Research Center. We are grateful for mice from Drs. Giorgio Trinchieri (NIH), Richard Flavell (Yale University), Roni Nowarski (Harvard University), and Kevin Barry (Fred Hutch) for the gift of mice, as well as G. Hill (Fred Hutch) and J. Trapani (Peter Mac Cancer Center, Australia) for helpful discussions. This research was supported by National Institutes of Health award numbers R01-HL145276 (J.A.D.), R01-HL165673 (J.A.D.), U01- AI70035 (J.A.D.), Project 1 of P01-AG052359 (J.A.D.), R35-HL171556 (J.A.D.) and the NCI Cancer Center Support Grant P30-CA015704. D.G. was supported by F30-HL165761. D.A. was supported by T32-GM007270.

## AUTHOR CONTRIBUTIONS

D.G. and J.A.D. conceived of the idea of this manuscript. D.G. and J.A.D. designed, analyzed, and interpreted experiments, and drafted the manuscript. K.C., D.A., M.W., L.I., P.D., E.L., S.S- S., S.K., and C.E. performed experiments and/or analyzed and helped interpret data. A.K., A.L., and M.V.D.B. aided in project discussion and sequencing data analysis. J.A.D. supervised the project.

## FIGURE LEGENDS

**Figure S1:**
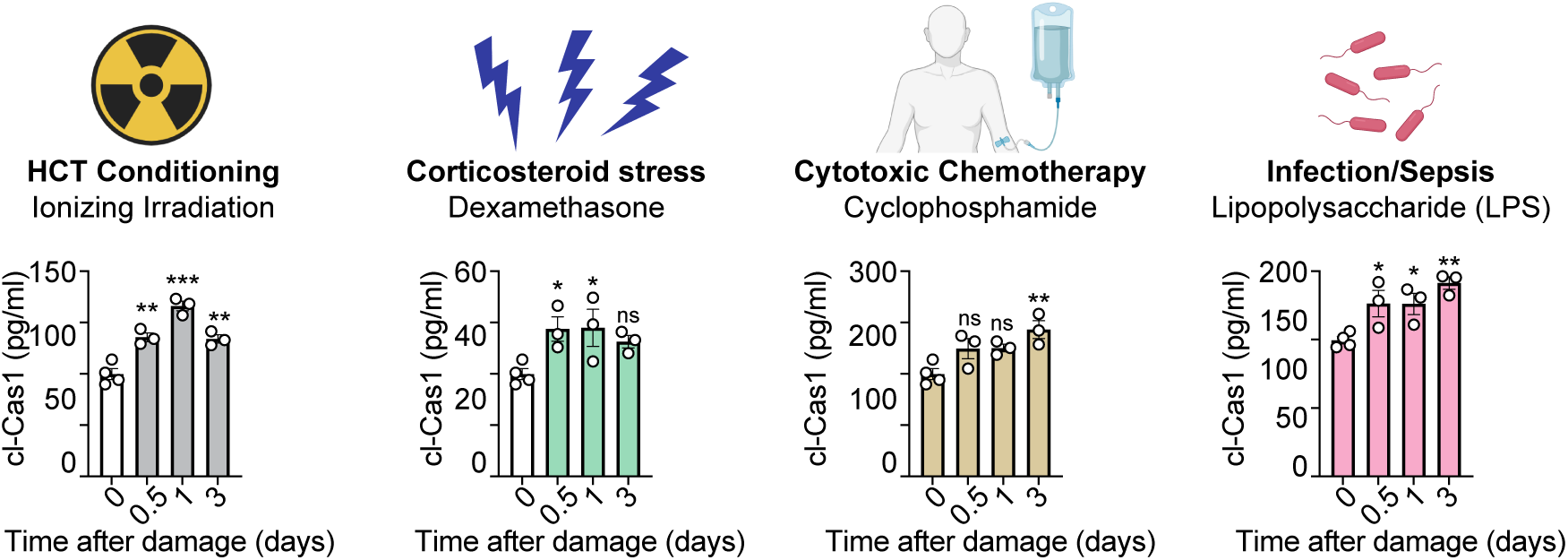
Cleaved Caspase 1 increases within the thymus following acute damages. Thymus supernatants generated from female 1-2 mo C57BL/6 mice given SL-TBI (550cGy), Dexamethasone (i.p., 20mg/kg), Cyclophosphamide (i.p., 200mg/kg) or LPS (i.p., 1.5mg/kg) and mature Caspase-1 assayed by ELISA on days 0, 0.5, 1 and 3 (n=3-4/group). Graphs represent mean ± SEM. *p<0.05, **p<0.01, ***p<0.001

**Figure S2:**
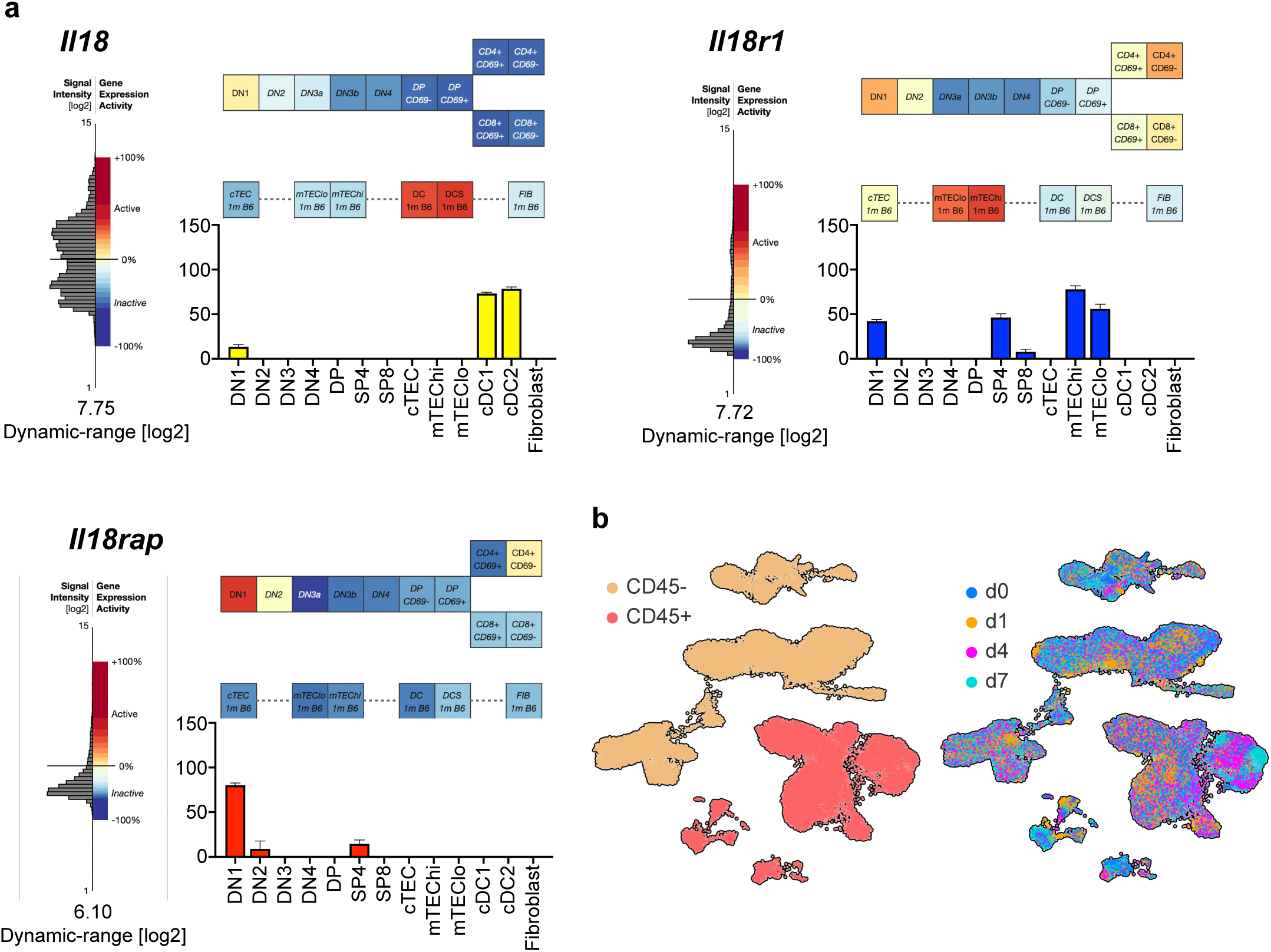
Cleaved Caspase 1 increases within the thymus following acute damages. **a,** Relative expression of *Il18*, *Il18r1* and *Il18rap* on subsets of thymus populations at baseline in 1mo mice. Data extracted from Gene Expression Commons using the “Complete thymocyte:stromal interaction model dataset” (https://gexc.riken.jp/models/475/) which is based on GSE56928^29^. **b**, UMAPs from Fig. 2A with color overlays indicating dataset (Left, RAG2- GFP-CD45+ or CD45-) or timepoint (right, day 0, 1, 4, 7 after SL-TBI).

**Figure S3:**
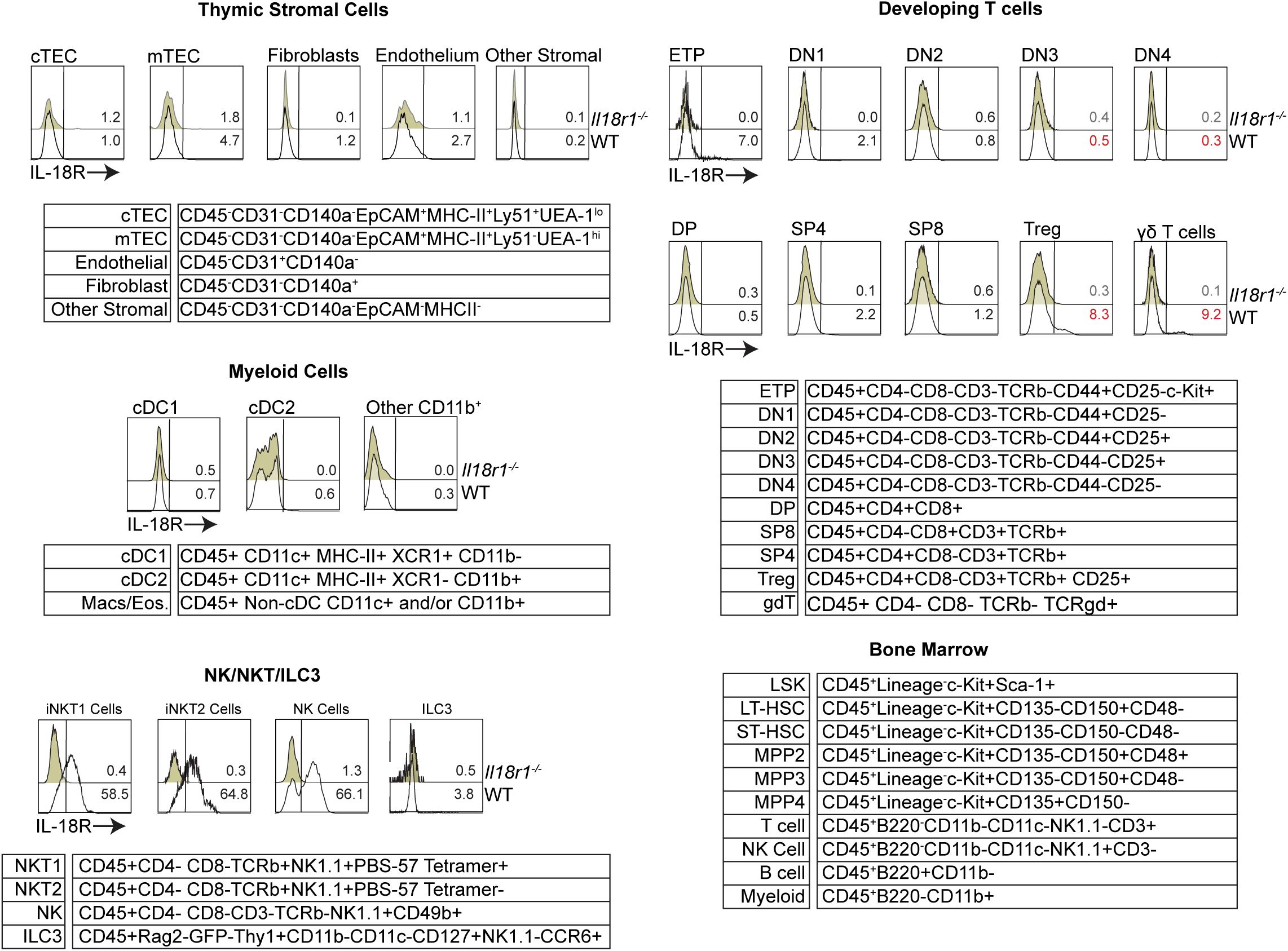
IL-18R expression on thymus populations. IL-18R expression on thymic cellular populations taken from female 1-2 mo C57BL/6 WT, Rag2gfp or Il18r1-/- mice (n=3-5/group). Tables show phenotypes used to gate individual populations.

**Figure S4:**
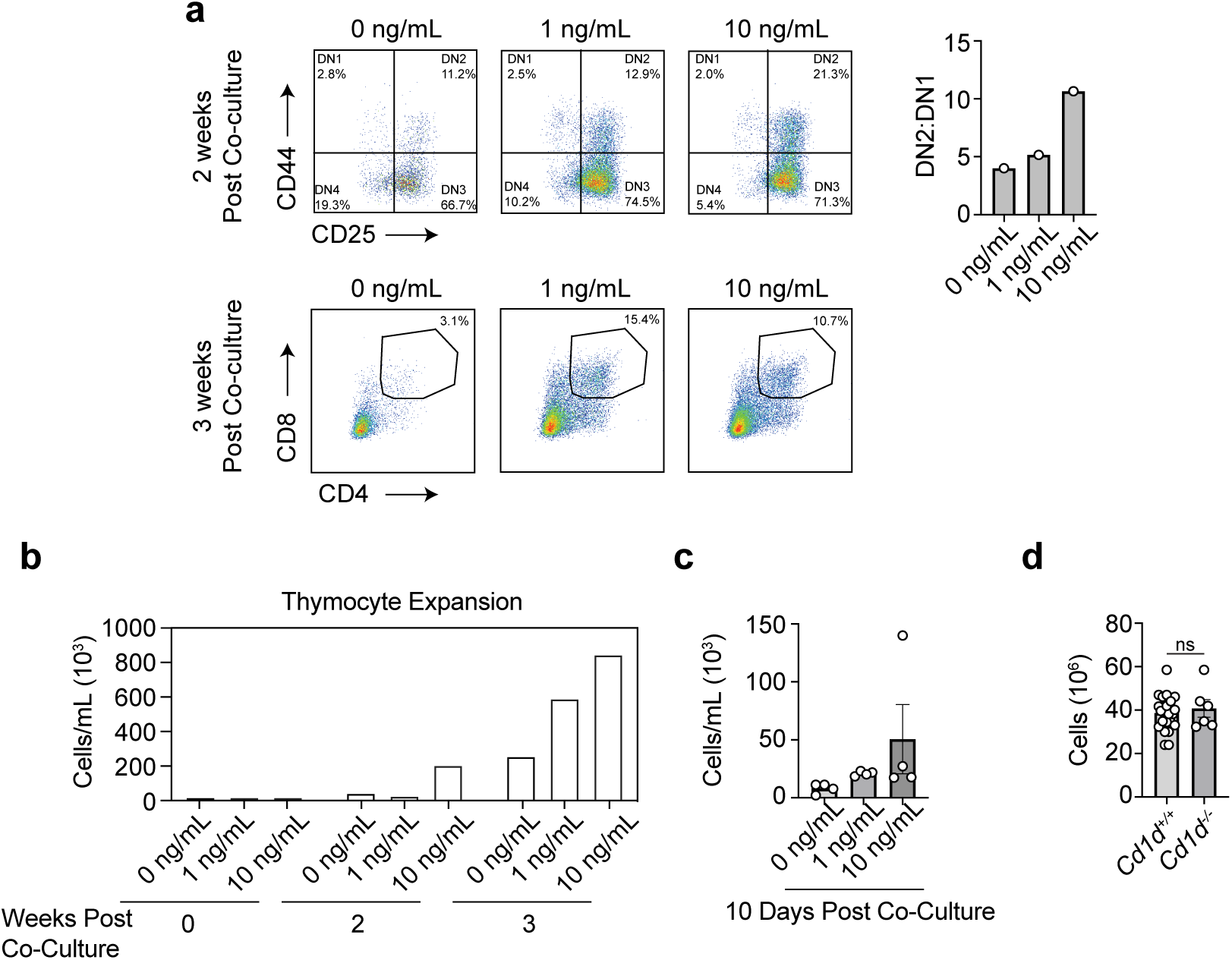
IL-18 promotes thymocyte expansion and differentiation *in vitro*. **a-b**, 50,000 lineage depleted bone marrow cells were co-cultured with OP9-DLL1^GFP^ fibroblasts confluent in a 6-well dish with 5ng/mL Flt3L plus 1ng/mL IL-7 and 0, 1 or 10 ng/mL rIL-18 (n=1/group). **a,** (Top) CD45^+^ Thy1^+^ CD4^-^ CD8^-^ DN1-4 thymocyte differentiation measured according to CD44 and CD25 expression 2 weeks post co-culture and (Bottom) CD45+Thy1+ thymocyte differentiation into CD4^+^ CD8^+^ Double Positive population 3 weeks post co-culture. (Right) Ratio of DN2:DN1 thymocyte differentiation 2 weeks post co-culture. **b,** Thymocyte expansion measured by non-adherent cell count quantified 2 and 3 weeks post co-culture with OP9-DLL1^GFP^ adherent cells. **c,** 50,000 bone marrow CD45+ Lineage- cKit+ Sca-1+ LSKs were FACS purified and co-cultured with OP9-DLL1^GFP^ fibroblasts confluent in a 6-well dish with 5ng/mL Ftl3L plus 1ng/mL IL-7 and 0, 1 or 10 ng/mL rIL-18 (n=4/group) and 10 days later, thymocyte expansion was quantified by measuring non-adherent cell expansion. **d,** Female 1-2 mo C57BL/6 WT (*Cd1d ^+/+^*) and *Cd1d ^-/-^* mice were given SL-TBI and thymus cellularity was measured 7 days later (n=7-8/group). Graphs represent mean ± SEM. *p<0.05, **p<0.01, ***p<0.001

